## Supplementary material for "NR2F2 regulation of interstitial to fetal Leydig cell differentiation in the testis: insights into differences of sex development": Suplemental figures and figure legends

**Supplementary figure legends**

**Figure S1: Cluster specific gene expression.** Feature plots of the normalized expression of cluster specific genes on the multiomic UMAP visualization of E14.5 testes.

**Figure S2: Interstitial clusters' specific gene expression.** (A) Feature plots of the normalized expression of interstitial clusters' specific genes on the multiomic UMAP visualization of E14.5 testes. (B) Immunofluorescence for cluster / cell type specific markers (Yellow) in E14.5 male testes. Samples were counterstained with DAPI (Grey). White arrowheads indicate immunopositive cells. White dashed box indicates the magnified area.

**Figure S3: Transcriptomic characterization of Leydig cells and Leydig progenitors.** Pathway analysis (Reactome 2022) of c0 (A) or c3 (B) DEGs. * indicates adjusted p-value <0.05.

**Figure S4: Lineage tracing of *Nr2f2* and *Wt1* positive cells in the fetal testis.**

(A) lineage tracing experiment of the fetal testis-derived *Nr2f2*+ cells in the *Nr2f2*-CreER; *CAG-Sun1/sfGFP* embryos. Cre activity was induced by tamoxifen administration at E14.5 and immunofluorescence was performed on E17.5 testes, for GFP (Green) and NR2F2 or NR5A1 (Magenta). White arrowheads indicate NR2F2 positive cells labelled with GFP. Yellow arrowheads indicate NR2F2 negative, GFP positive or GFP NR5A1 double positive Leydig cells. (B) Lineage tracing of the fetal testis-derived Wt1+ cells in the *Wt1*-CreER; *CAG-*

*Sun1/sfGFP* embryos was induced by tamoxifen administration at E11.5 and E12.5. Immunofluorescence for GFP (Green) and NR2F2 (Magenta) was performed on E14.5 or E17.5 testes. White dashed box indicates the magnified area. Image created with BioRender.com

**Figure S5: Characterization of the *Nr2f2* knockout testicular phenotype.** Pathway analysis (Reactome 2022) of all the downregulated (A) or all the upregulated (B) genes in the knockout testes vs the control. * indicates adjusted p-value <0.05. (C) Quantification of the gonadal area in control vs *Nr2f2* conditional knockout gonads. Gonadal area of each sample was normalized to the average area of the control testes. (D) Quantification of the % of interstitial area in control and *Nr2f2* cKO gonads.

**Figure S6: Bulk RNA-seq deconvolution using single-nuclei RNA-seq data.** Bulk RNA-seq deconvolution was performed using our E14.5 testicular single-nuclei data. Heatmaps were generated using the average expression for each cluster of the upregulated (A) or downregulated (B) genes in *Nr2f2* knockout. Genes were then hierarchical grouped (grp#) based on the expression patterns across the 12 clusters. Gene ontology (biological process, GO:BP) analysis was performed on each of the different gene groups and the most representative statistically significant (adj P value<0.05) is presented next to each gene group.

**Figure S7: Pathway analysis of the ChIP-seq and ATAC-seq data.** (A) Cell cycle status (G1, G2M and S) of each cell per single-nuclei multiomic cluster, represented as a % of the total number of cells per each cluster. Pathway analysis (WikiPathway 2023) of all NR2F2 target genes (B) or all NR2F2 TSS proximal target genes (C). Pathway analysis (Reactome 2022) of all C0 (D) or C3 (E) DEGs with a significant NR2F2 peak associated. Pathway analysis (Reactome 2022) of all the linked genes to a differentially accessible peak in c0 (F) or c3 (G). (H) Pathway analysis (Reactome 2022) of all differentially accessible peaks on c0 vs c3, containing an NR2F2 ChIP-seq peak. * Indicates adjusted p-value <0.05.

**Supplementary table legends**

**Table S1.** Primary and secondary antibodies used for immunofluorescence.

**Table S2.** Differentially expressed genes per single-nuclei multiomic cluster from E14.5 testes.

**Table S3.** Differentially expressed genes between c0 (interstitial cells) and c3 (leydig cells). This table also contains the pathway analysis (Reactome 2022) for c0 and c3 DEGs.

**Table S4.** Bulk RNA-seq data from E14.5 control and *Nr2f2* conditional knockout testes. This table contains QC summary, fragment size plot, fragment size data and the differential expression analysis. This table also contains the pathway analysis (Reactome 2022) for upregulated and downregulated genes in *Nr2f2* cKO vs control.

**Table S5.** List of genes per gene group from the bulk RNA-seq deconvolution of the upregulated and downregulated DEGs between E14.5 *Nr2f2* cKO and control testes. This table contains gene ontology analysis (biological process) for each of the different gene groups in the upregulated and downregulated datasets.

**Table S6.** Statistically significant NR2F2 peaks called in E14.5 testes. This table includes the genomic context, the closest gene and the distance in bp to that gene. This table also contains the pathway analysis (Wikipathway 2023) for all or TSS proximal NR2F2 ChIP-seq targets and for all the NR2F2 ChIP-seq targets that are c0 and c3 DEGs (Reactome 2022).

**Table S7.** Differentially accessible peaks between c0 (interstitial cells) and c3 (Leydig cells). This table includes the ATAC peak range, the ATAC peak, the distance to the nearest NR2F2 ChIP-seq peak and the linked genes to the peak. This table also contains the pathway analysis (Reactome 2022) for all the genes linked to c0 and c3 specific peaks, and the genes linked to c0 specific peaks that also contains a NR2F2 ChIP-seq binding peak.

**Table S8.** Motifs enriched in differentially accessible peaks in c0 and c3. This table includes the list of enriched motifs for c0 and c3, the DEGs analysis of the motifs in c0 and c3 and the combination analysis with the *Nr2f2* knockout RNA-seq and the NR2F2 ChIP-seq. This table also contains the Motifs enriched in differentially accessible peaks in c0 vs c3 that that also contains a NR2F2 ChIP-seq binding peak.

### Figure S1

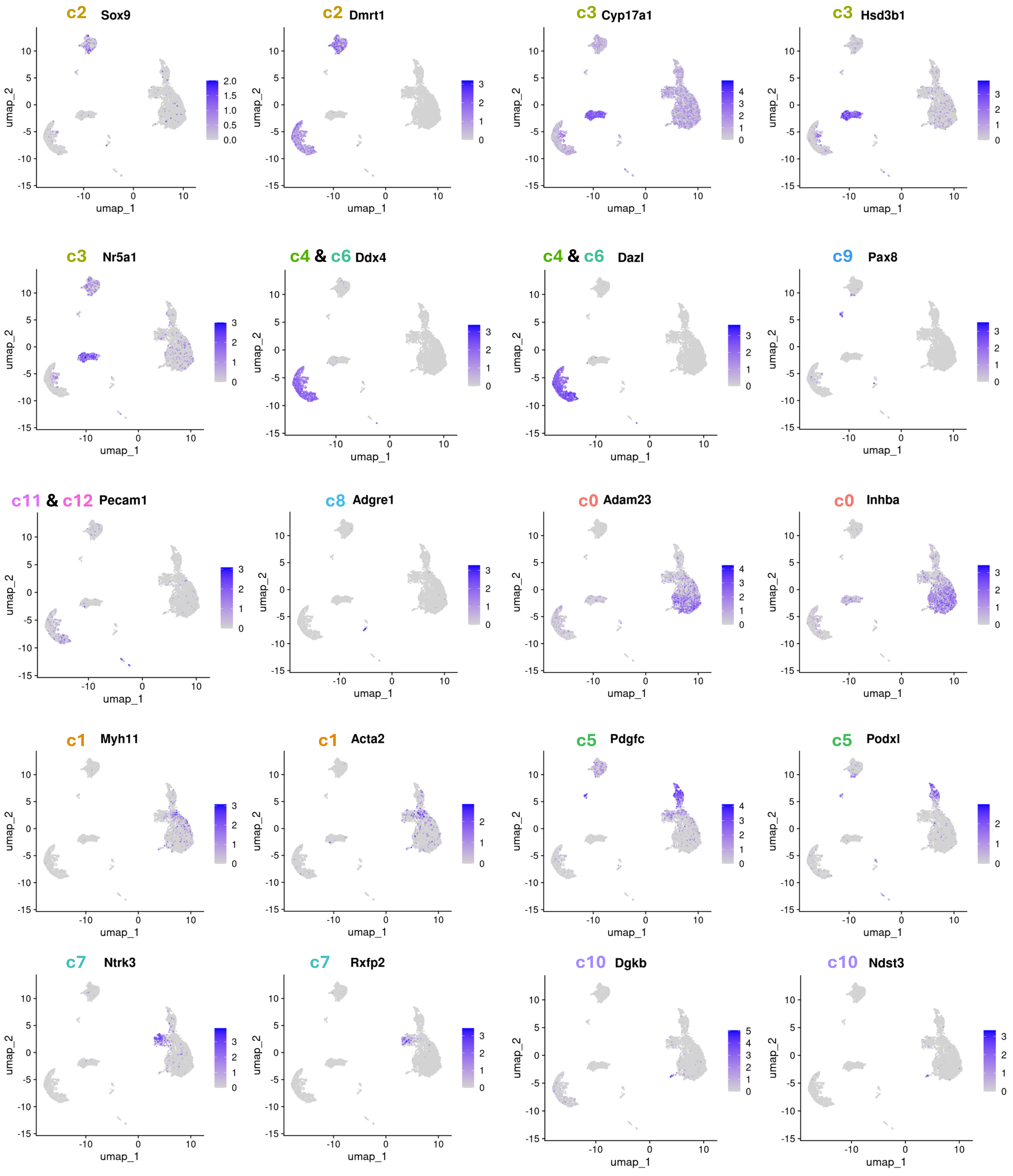

c2 Sox9
c2 Dmrt1
c3 Cyp17a1
c3 Hsd3b1
c3 Nr5a1
c4 & c6 Ddx4
c4 & c6 Dazl
c9 Pax8
c11 & c12 Pecam1
c8 Adgre1
c0 Adam23
c0 Inhba
c1 Myh11
c1 Acta2
c5 Pdgfc
c5 Podxl
c7 Ntrk3
c7 Rxfp2
c10 Dgkb
c10 Ndst3
umap_1
umap_2

### Figure S2

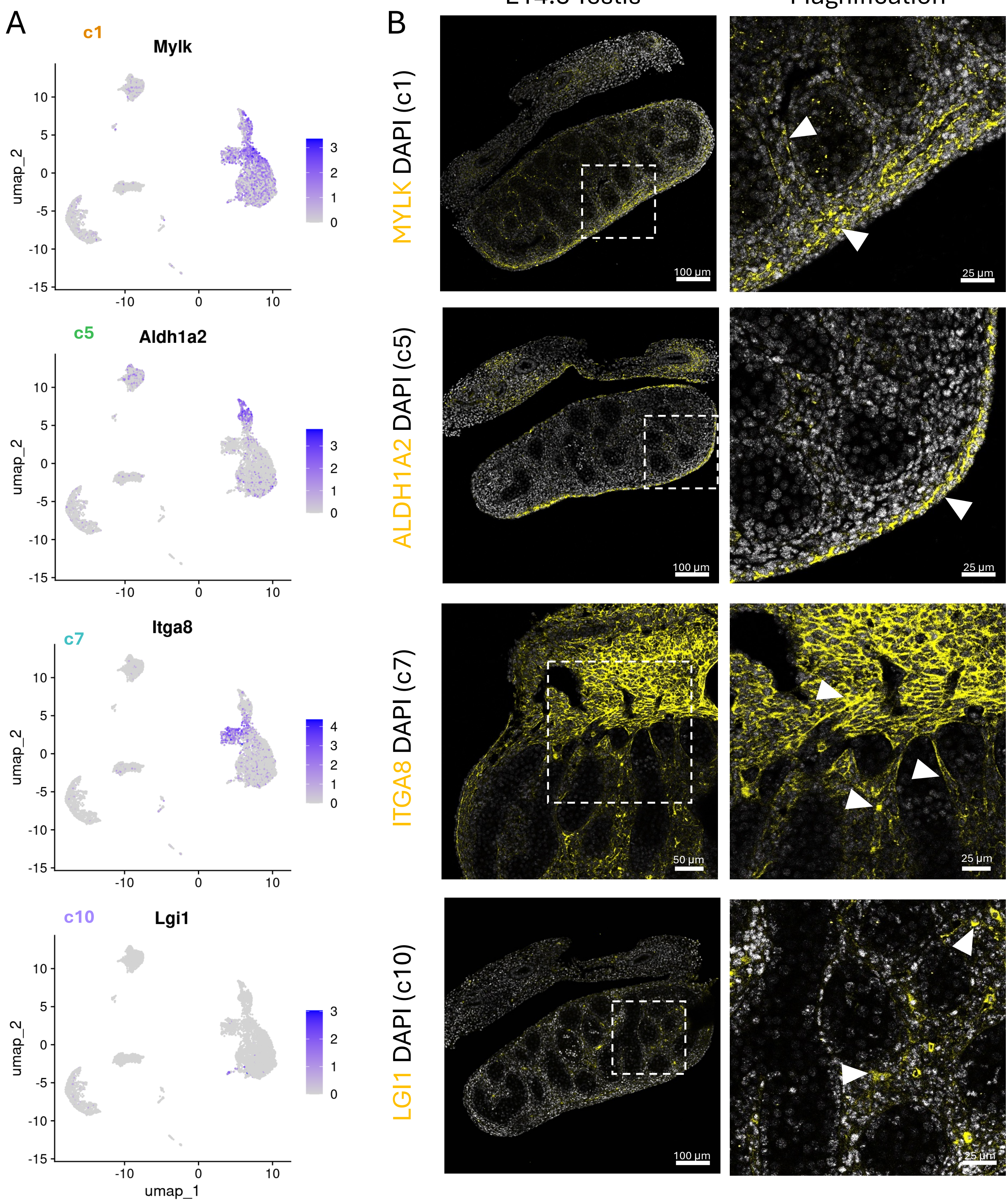

A
c1
Mylk
c5
Aldh1a2
c7
Itga8
c10
Lgi1
umap_2
umap_1
B
E14.5 Testis
Magnification
MYLK DAPI (c1)
ALDH1A2 DAPI (c5)
ITGA8 DAPI (c7)
LGI1 DAPI (c10)
100 µm
50 µm
25 µm

### Figure S3

c0 Interstitial

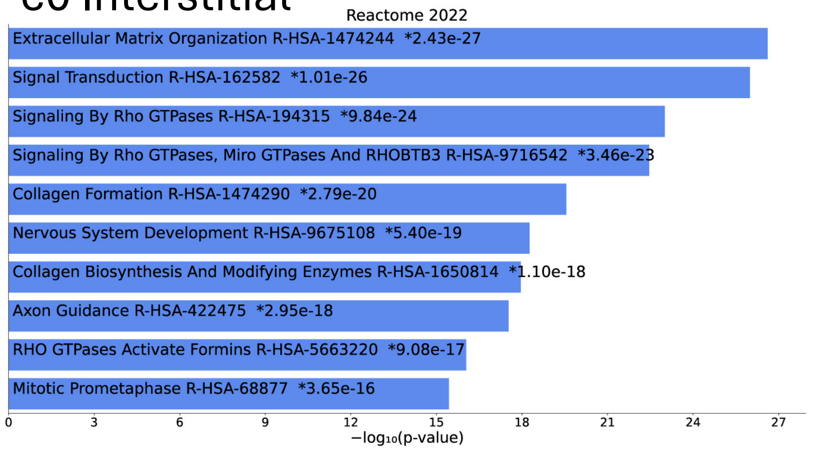


c3 Leydig

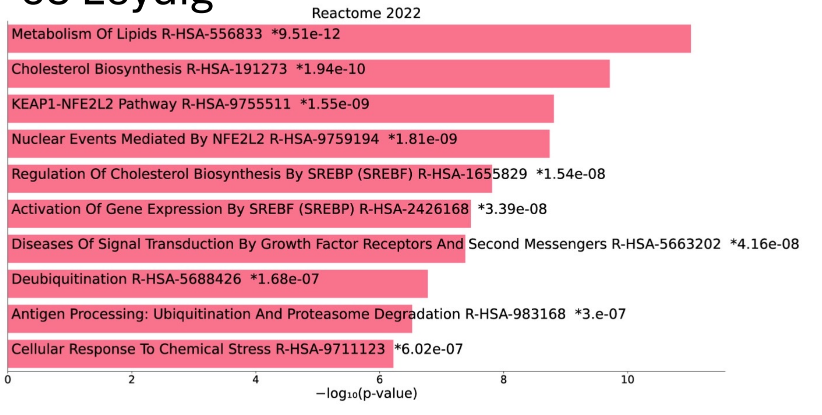

Figure S4

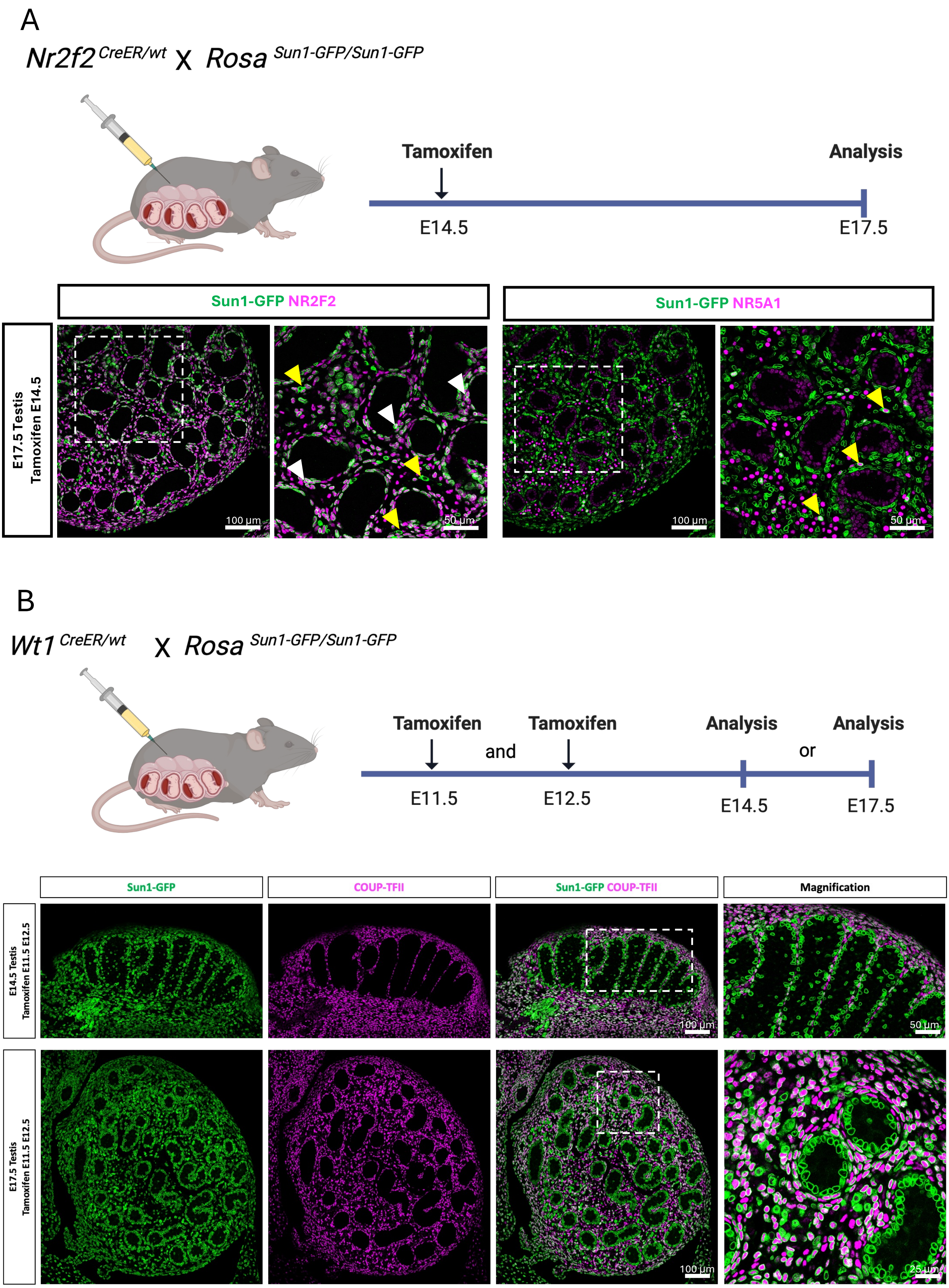
A
Nr2f2 CreER/wt X Rosa Sun1-GFP/Sun1-GFP
Tamoxifen
Analysis
E14.5
E17.5
Sun1-GFP NR2F2
Sun1-GFP NR5A1
E17.5 Testis
Tamoxifen E14.5
100 µm
50 µm
100 µm
50 µm
B
Wt1 CreER/wt X Rosa Sun1-GFP/Sun1-GFP
Tamoxifen
and
Tamoxifen
Analysis
or
Analysis
E11.5
E12.5
E14.5
E17.5
Sun1-GFP
COUP-TFII
Sun1-GFP COUP-TFII
Magnification
E14.5 Testis
Tamoxifen E11.5 E12.5
100 µm
50 µm
E17.5 Testis
Tamoxifen E11.5 E12.5
100 µm
25 µm

Figure S5:

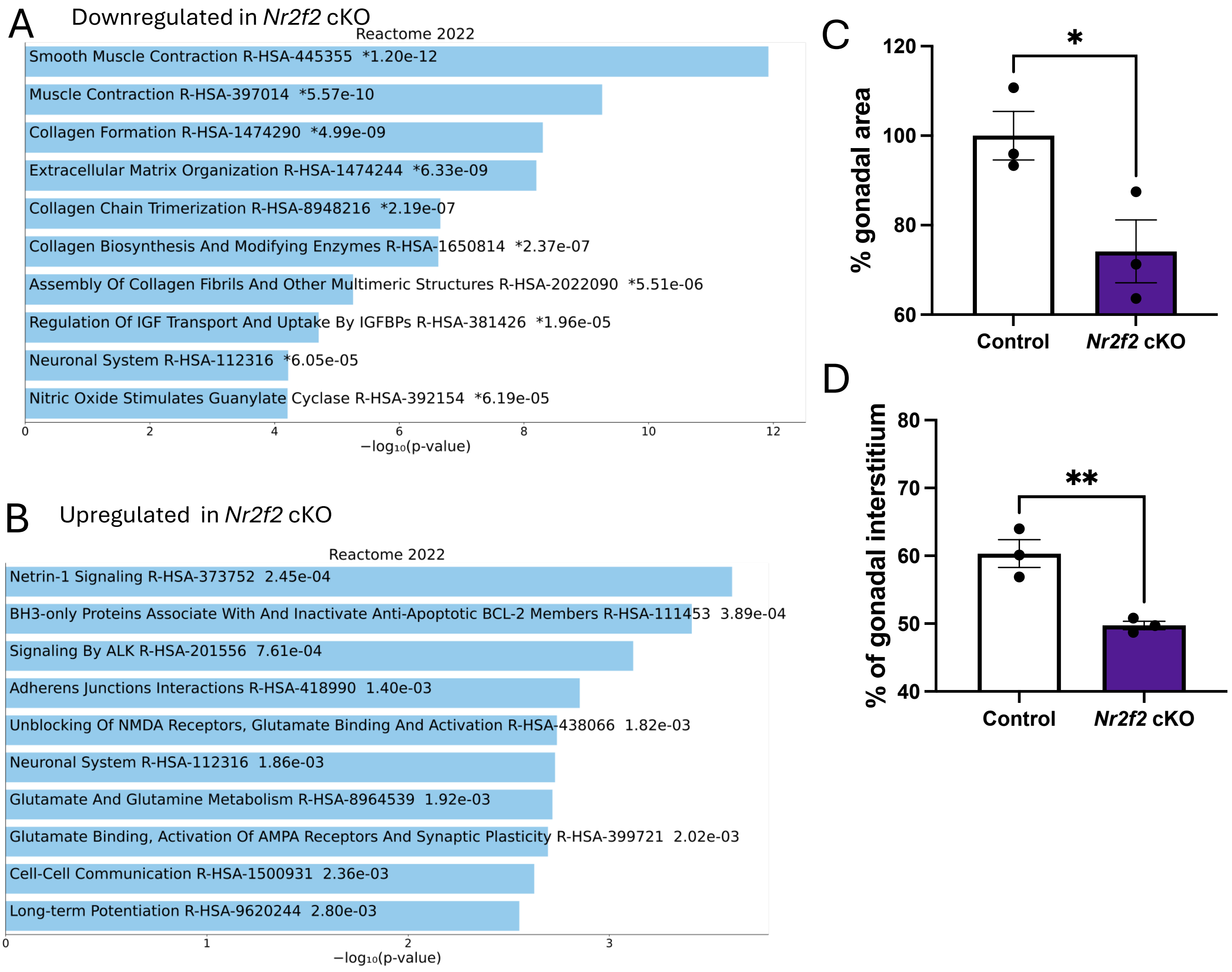
A
Downregulated in Nr2f2 cKO
Reactome 2022
Smooth Muscle Contraction R-HSA-445355 *1.20e-12
Muscle Contraction R-HSA-397014 *5.57e-10
Collagen Formation R-HSA-1474290 *4.99e-09
Extracellular Matrix Organization R-HSA-1474244 *6.33e-09
Collagen Chain Trimerization R-HSA-8948216 *2.19e-07
Collagen Biosynthesis And Modifying Enzymes R-HSA-1650814 *2.37e-07
Assembly Of Collagen Fibrils And Other Multimeric Structures R-HSA-2022090 *5.51e-06
Regulation Of IGF Transport And Uptake By IGFBPs R-HSA-381426 *1.96e-05
Neuronal System R-HSA-112316 *6.05e-05
Nitric Oxide Stimulates Guanylate Cyclase R-HSA-392154 *6.19e-05
0
2
4
6
8
10
12
−log10(p-value)
B
Upregulated in Nr2f2 cKO
Reactome 2022
Netrin-1 Signaling R-HSA-373752 2.45e-04
BH3-only Proteins Associate With And Inactivate Anti-Apoptotic BCL-2 Members R-HSA-111453 3.89e-04
Signaling By ALK R-HSA-201556 7.61e-04
Adherens Junctions Interactions R-HSA-418990 1.40e-03
Unblocking Of NMDA Receptors, Glutamate Binding And Activation R-HSA-438066 1.82e-03
Neuronal System R-HSA-112316 1.86e-03
Glutamate And Glutamine Metabolism R-HSA-8964539 1.92e-03
Glutamate Binding, Activation Of AMPA Receptors And Synaptic Plasticity R-HSA-399721 2.02e-03
Cell-Cell Communication R-HSA-1500931 2.36e-03
Long-term Potentiation R-HSA-9620244 2.80e-03
0
1
2
3
−log10(p-value)
C
% gonadal area
120
100
80
60
*
Control
Nr2f2 cKO
D
% of gonadal interstitium
80
70
60
50
40
**
Control
Nr2f2 cKO

### Figure S6:

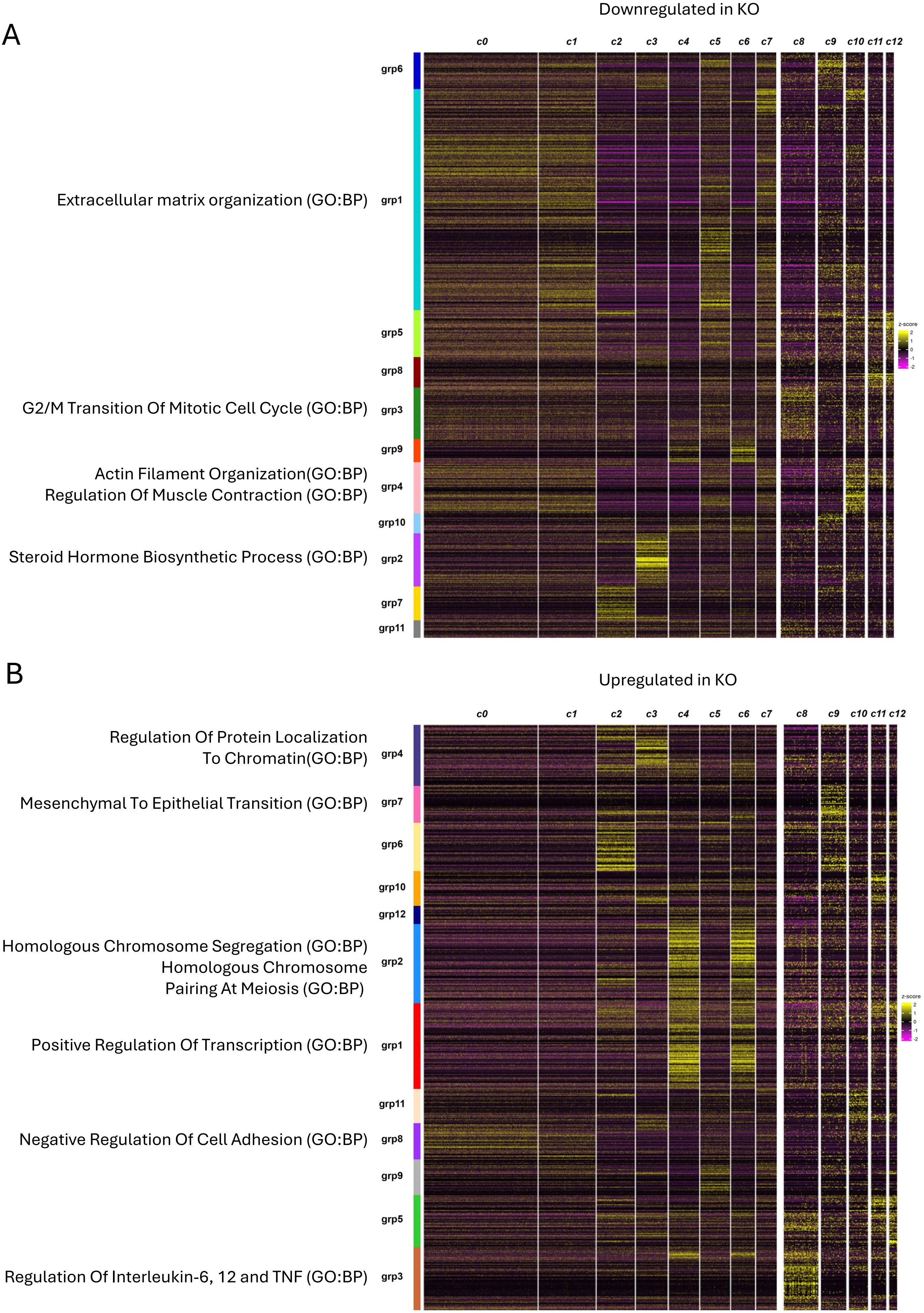
A
Downregulated in KO
c0
c1
c2
c3
c4
c5
c6
c7
c8
c9
c10
c11
c12
grp6
Extracellular matrix organization (GO:BP)
grp1
grp5
grp8
G2/M Transition Of Mitotic Cell Cycle (GO:BP)
grp3
grp9
Actin Filament Organization(GO:BP)
Regulation Of Muscle Contraction (GO:BP)
grp4
grp10
Steroid Hormone Biosynthetic Process (GO:BP)
grp2
grp7
grp11
z-score
B
Upregulated in KO
Regulation Of Protein Localization
To Chromatin(GO:BP)
grp4
Mesenchymal To Epithelial Transition (GO:BP)
grp7
grp6
grp10
grp12
Homologous Chromosome Segregation (GO:BP)
Homologous Chromosome
Pairing At Meiosis (GO:BP)
grp2
Positive Regulation Of Transcription (GO:BP)
grp1
grp11
Negative Regulation Of Cell Adhesion (GO:BP)
grp8
grp9
grp5
Regulation Of Interleukin-6, 12 and TNF (GO:BP)
grp3

### Figure S7:

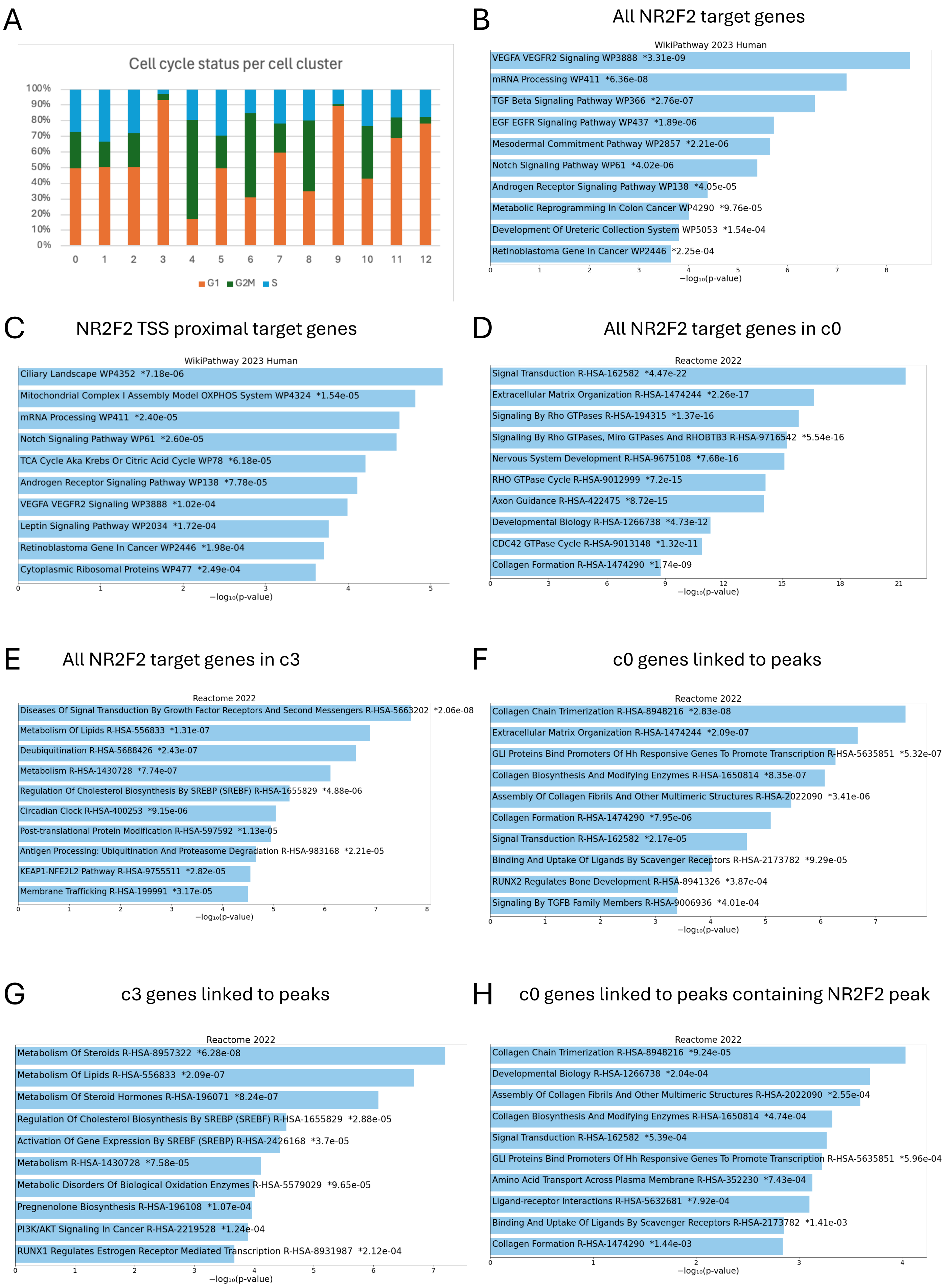
A
Cell cycle status per cell cluster
100%
90%
80%
70%
60%
50%
40%
30%
20%
10%
0%
0 1 2 3 4 5 6 7 8 9 10 11 12
G1 G2M S
B
All NR2F2 target genes
WikiPathway 2023 Human
VEGFA VEGFR2 Signaling WP3888 *3.31e-09
mRNA Processing WP411 *6.36e-08
TGF Beta Signaling Pathway WP366 *2.76e-07
EGF EGFR Signaling Pathway WP437 *1.89e-06
Mesodermal Commitment Pathway WP2857 *2.21e-06
Notch Signaling Pathway WP61 *4.02e-06
Androgen Receptor Signaling Pathway WP138 *4.05e-05
Metabolic Reprogramming In Colon Cancer WP4290 *9.76e-05
Development Of Ureteric Collection System WP5053 *1.54e-04
Retinoblastoma Gene In Cancer WP2446 *2.25e-04
0 1 2 3 4 5 6 7 8
−log₁₀(p-value)
C
NR2F2 TSS proximal target genes
WikiPathway 2023 Human
Ciliary Landscape WP4352 *7.18e-06
Mitochondrial Complex I Assembly Model OXPHOS System WP4324 *1.54e-05
mRNA Processing WP411 *2.40e-05
Notch Signaling Pathway WP61 *2.60e-05
TCA Cycle Aka Krebs Or Citric Acid Cycle WP78 *6.18e-05
Androgen Receptor Signaling Pathway WP138 *7.78e-05
VEGFA VEGFR2 Signaling WP3888 *1.02e-04
Leptin Signaling Pathway WP2034 *1.72e-04
Retinoblastoma Gene In Cancer WP2446 *1.98e-04
Cytoplasmic Ribosomal Proteins WP477 *2.49e-04
0 1 2 3 4 5
−log₁₀(p-value)
D
All NR2F2 target genes in c0
Reactome 2022
Signal Transduction R-HSA-162582 *4.47e-22
Extracellular Matrix Organization R-HSA-1474244 *2.26e-17
Signaling By Rho GTPases R-HSA-194315 *1.37e-16
Signaling By Rho GTPases, Miro GTPases And RHOBTB3 R-HSA-9716542 *5.54e-16
Nervous System Development R-HSA-9675108 *7.68e-16
RHO GTPase Cycle R-HSA-9012999 *7.2e-15
Axon Guidance R-HSA-422475 *8.72e-15
Developmental Biology R-HSA-1266738 *4.73e-12
CDC42 GTPase Cycle R-HSA-9013148 *1.32e-11
Collagen Formation R-HSA-1474290 *1.74e-09
0 3 6 9 12 15 18 21
−log₁₀(p-value)
E
All NR2F2 target genes in c3
Reactome 2022
Diseases Of Signal Transduction By Growth Factor Receptors And Second Messengers R-HSA-5663202 *2.06e-08
Metabolism Of Lipids R-HSA-556833 *1.31e-07
Deubiquitination R-HSA-5688426 *2.43e-07
Metabolism R-HSA-1430728 *7.74e-07
Regulation Of Cholesterol Biosynthesis By SREBP (SREBF) R-HSA-1655829 *4.88e-06
Circadian Clock R-HSA-400253 *9.15e-06
Post-translational Protein Modification R-HSA-597592 *1.13e-05
Antigen Processing: Ubiquitination And Proteasome Degradation R-HSA-983168 *2.21e-05
KEAP1-NFE2L2 Pathway R-HSA-9755511 *2.82e-05
Membrane Trafficking R-HSA-199991 *3.17e-05
0 1 2 3 4 5 6 7 8
−log₁₀(p-value)
F
c0 genes linked to peaks
Reactome 2022
Collagen Chain Trimerization R-HSA-8948216 *2.83e-08
Extracellular Matrix Organization R-HSA-1474244 *2.09e-07
GLI Proteins Bind Promoters Of Hh Responsive Genes To Promote Transcription R-HSA-5635851 *5.32e-07
Collagen Biosynthesis And Modifying Enzymes R-HSA-1650814 *8.35e-07
Assembly Of Collagen Fibrils And Other Multimeric Structures R-HSA-2022090 *3.41e-06
Collagen Formation R-HSA-1474290 *7.95e-06
Signal Transduction R-HSA-162582 *2.17e-05
Binding And Uptake Of Ligands By Scavenger Receptors R-HSA-2173782 *9.29e-05
RUNX2 Regulates Bone Development R-HSA-8941326 *3.87e-04
Signaling By TGFB Family Members R-HSA-9006936 *4.01e-04
0 1 2 3 4 5 6 7
−log₁₀(p-value)
G
c3 genes linked to peaks
Reactome 2022
Metabolism Of Steroids R-HSA-8957322 *6.28e-08
Metabolism Of Lipids R-HSA-556833 *2.09e-07
Metabolism Of Steroid Hormones R-HSA-196071 *8.24e-07
Regulation Of Cholesterol Biosynthesis By SREBP (SREBF) R-HSA-1655829 *2.88e-05
Activation Of Gene Expression By SREBF (SREBP) R-HSA-2426168 *3.7e-05
Metabolism R-HSA-1430728 *7.58e-05
Metabolic Disorders Of Biological Oxidation Enzymes R-HSA-5579029 *9.65e-05
Pregnenolone Biosynthesis R-HSA-196108 *1.07e-04
PI3K/AKT Signaling In Cancer R-HSA-2219528 *1.24e-04
RUNX1 Regulates Estrogen Receptor Mediated Transcription R-HSA-8931987 *2.12e-04
0 1 2 3 4 5 6 7
−log₁₀(p-value)
H
c0 genes linked to peaks containing NR2F2 peak
Reactome 2022
Collagen Chain Trimerization R-HSA-8948216 *9.24e-05
Developmental Biology R-HSA-1266738 *2.04e-04
Assembly Of Collagen Fibrils And Other Multimeric Structures R-HSA-2022090 *2.55e-04
Collagen Biosynthesis And Modifying Enzymes R-HSA-1650814 *4.74e-04
Signal Transduction R-HSA-162582 *5.39e-04
GLI Proteins Bind Promoters Of Hh Responsive Genes To Promote Transcription R-HSA-5635851 *5.96e-04
Amino Acid Transport Across Plasma Membrane R-HSA-352230 *7.43e-04
Ligand-receptor Interactions R-HSA-5632681 *7.92e-04
Binding And Uptake Of Ligands By Scavenger Receptors R-HSA-2173782 *1.41e-03
Collagen Formation R-HSA-1474290 *1.44e-03
0 1 2 3 4
−log₁₀(p-value)
